# Identification of Evolutionary Trade-Offs Associated with High-Level Colistin Resistance in *Acinetobacter baumannii*

**DOI:** 10.1101/2025.01.17.633615

**Authors:** Kusumita Acharya, Upasana Bhattacharya, Shatarupa Biswas, Mallika Ghosh, Arijit Bhattacharya

## Abstract

Colistin (COL) belongs to the polymyxin group of drugs which possesses a positive charge and interacts with lipopolysaccharide (LPS) of Gram-negative bacterial outer membrane. Additionally, it can penetrate the cell membrane and disrupt the integrity of the phospholipid leading to cell death. Resistance against COL has been reported in *Acinetobacter baumannii*, a bacteria in the ‘ESKAPE’ group of ‘priority pathogens’ of the World Health Organization. Multiple and pan-drug-resistant strains of *A. baumannii* are rising in many medical facilities and they are becoming resistant to last-resort antibiotics including colistin. Though plasmid-encoded acquisition of *mcr*-1 has been associated with clinical resistance, drug efflux, complete loss of LPS by inactivation of the biosynthetic pathway (*lpx*ACD), and modifications of target LPS by-products of chromosomal *pmr*CAB genes has been ascribed with resistance evolution against COL. To further delve into the evolutionary dynamics of COL-resistance here we report experimental evolution of extreme COL-resistance by adaptive evolution of the reference strain *A. baumannii ATCC*19606. Phenotypic characterization has been performed for the evolved strains, which demonstrated hyperbiofilm phenotype and striking decrease in fitness. A comprehensive antibiotic susceptibility profiling has been performed on the evolved strains. Whole genome sequencing of the resistant strains led to the identification of mutations prospectively associated with COL resistance. Phenotypic characterization of three COL-resistant clinical isolates of *A. baumannii* revealed similarity with experimentally evolved resistant mutants at least in one of the isolates.

## Introduction

Bacterial Antimicrobial resistance (AMR) is arguably the major challenge in the global health scenario in the present century with approximately 4.71 million deaths associated with it in 2021 with an immense rise in Gram-negative infection burden [1]. Among the major Gram-negative pathogens, *Acinetobacter baumannii* (Ab) has emerged as a critical concern as a nosocomial opportunistic pathogen due to its global dissemination and acquisition of resistance against last-resort antibiotics [2]. The bacteria cause infections in the respiratory tract, urinary tract, other soft tissues, and wounds with an attributable mortality rate nearing 35% [3]. The bacterium demonstrates considerable phenotypic and genotypic diversity with a four times higher propensity of multidrug resistance (MDR) compared to other nosocomial pathogens and rapid development of pan-drug resistance [4]. Though carbapenem-resistant Ab (CRAB) has been highlighted as a ‘priority pathogen’ with an immediate need for new antibiotics (https://www.who.int/publications/i/item/9789240093461), reports for clinical resistance against the other last-resort antibiotic colistin are increasing owing to increased use of the antibiotic to treat infections caused by MDR isolates [5].

The bactericidal antibiotic colistin (polymyxin E, COL) is a polycationic lipopeptide, comprising a cyclic decapeptide conjugated to a fatty acyl chain [6]. The lipopeptide moiety can directly bind to the negatively charged phosphates of lipid-A part of lipopolysaccharide, a major constituent of the outer membrane (OM) of Gram-negative bacteria. The interaction displaces Ca^2+^ and Mg^2+^ ions from lipid-A and impairs the integrity of OM. Following its passage to the cytoplasmic membrane it manifests leakage of cellular contents [7]. In addition to the well-studied mechanism, COL induces the production of hydroxyl radicals, contributing to immediate cell death. This hydroxyl radical-mediated cell death pathway is observed in clinical Ab isolates [8,9].

Among diverse groups of Gram-negative bacteria, COL-resistance is conferred by chromosomal mutations in LPS biosynthesizing genes and genes encoding two-component systems modulating LPS biosynthesis, efflux pumps, and plasmid-encoded genes belonging to phosphoethanolamine transferase family [10]. In *A. baumannii*, COL-resistance has been studied with resistant experimentally raised mutants and resistant clinical isolates [7]. Mutations in LPS core biosynthesizing genes like *lps*B have been attributed to the high level of intrinsic COL-resistance [11]. In line with gene inactivation identified in other bacteria, nucleotide substitution, insertion-deletion (InDels), and insertion inactivation of lipid-A biosynthetic genes *lpx*A, *lpx*C, and *lpx*D have been identified in several LPS deficient COL^R^ mutants [12–14]. Deletion in the translocase LptD encoding gene was identified to be associated with COL-resistance in Ab ATCC 19606, where LPS biosynthesis is not essential *in vitro* [15]. Modification of lipid-A by addition of phosphoethanolamine by phosphoethanolamine transferase PmrC mitigates COL binding to OM [16]. PmrC expression is controlled by the two-component system containing the response regulator PmrA and the sensor histidine kinase PmrB [17,18]. Mutations in both *pmr*A and *pmr*B and overexpression as well, have strongly been associated with various degrees of COL-resistance [5,19]. The plasmid harbored mobilized COL resistance genes (*mcr*) that have been identified in clinical isolates of Ab. The gene is transmitted horizontally and encodes a phosphoethanolamine transferase (PET), a lipid modifier [20]. Of the ten different *mcr* genes and various variants identified, *mcr1* and *mcr4*.3 have been commonly identified in COL^R^ isolates of Ab [21,22]. COL exposure to Ab also inflicts ‘colistin dependency’ for *in vitro* growth [23]. Apart from these mechanisms, Ab can develop resistance through the efflux system EmrAB, which pumps out antibiotics, and the production of enzymes facilitating proteolytic cleavage of drugs [24,25].

Succeeding previous studies to characterize COL-resistance in Ab, in this study, extremely COL-resistant *A. baumannii* ATCC19606 (Ab19606) mutants evolved experimentally by step-wise selection. The mutants evolved are compromised in terms of fitness and are profound biofilm formers. Albeit no *mcr* or change in copy number for *pmr*A or *pmr*B could be detected, the mutants are impaired in outer membrane integrity. Whole genome sequencing identified an intentional mutation leading to the duplication of proline in LpxA2, along with mutations in several other genes. Comprehensive cross-resistance and collateral sensitivity (CR-CS) profiling revealed sensitivity to several antibiotics as possible evolutionary trade-offs including susceptibility to vancomycin (VAN), which corroborated earlier reports [26]. Three COL-R clinical isolates of Ab were also characterized, one of which indicated hyperbiofilm formation and antibiotic sensitivity trade-offs similar to the laboratory-evolved mutants.

## Methodology

### Strains and reagents

*Acinetobacter baumannii* (ATCC19606) (Ab19606) was maintained in Luria-Bertani agar and broth (Himedia). LB broth with 1% glucose is used for biofilm assays. Leeds Acinetobacter Agar (Himedia) was used for selective assays involving Ab. Antimicrobial susceptibility tests were performed in Mueller Hinton Broth (Himedia). All the reagents including antibiotics like colistin (COL) and ciprofloxacin (CIP) were purchased from Sigma-Aldrich unless mentioned otherwise. Antibiotic disks and antibiotic strips were procured from Himedia.

### Laboratory evolution of resistant mutants

A source clonal population was generated from a single clone of Ab19606. Laboratory evolution experiments involve serial transfer of growing bacterial populations in parallel for hundreds of generations with gradually increasing concentrations of each of COL and CIP. For each step of adaptation, cultures were allowed to grow till OD_600 nm_ of 0.8 (late log phase). In parallel, untreated cultures were passaged as control lines (Fig. S1). Clones derived from single colonies derived from evolved lines were verified by PCR amplification of genomic DNA for *gapdh* and 16S rRNA gene using primer pairs -AbgapdhF: ATGCAACGTATCGCCATT, AbgapdhR: TCGTACATGACACACTCGAT and Ab16SF: GAATAAGCACCGGCTAACTCTGT and Ab16SR: TAAGGTTCTTCGCGTTGCAT using T100 Thermal Cycler (Biorad), followed by Sanger sequencing. The clones were also profiled for changes in susceptibilities against COL and CIP.

### Obtaining clinical isolates

Three clinical isolates were obtained with identifiable COL-resistance. Pathological “Resistance (R)” was assigned according to automated AST assay with VITEK 2 system version 9.02 in accordance with Global CLSI-based 2023. After collection, strain confirmations were done on Leeds *Acinetobacter* agar.

### Characterizing clonal population of evolved lines and clinical isolates

The level of resistance of each isolate was tested by determining MIC through microdilution as described earlier [27]. Alongside MBC each isolate was evaluated by determining CFU at break point concentrations. The extent of resistance was expressed in terms of fold-MIC change compared to control lines. MICs were determined as suggested by the Clinical and Laboratory Standards Institute (CLSI) guidelines with a cell number of 5X10^5^/ ml using 9-step serial dilutions in 96-well tissue culture plates. For measurement of small changes, 1.2–1.5 times dilutions were used in macrodilution.

### Growth and fitness assessment of the resistant clonal population against source clonal population

The fitness cost of the resistant mutations was analyzed by comparative growth profiling as described by Lenski et al. [28] where the growth parameter for a strain is the number of doublings that it experiences over a given period of time.

### Screening for cross-resistance and collateral sensitivity

*Disk-diffusion susceptibility testing-* 0.5 McFarland matched bacterial suspension was spread on LB plates and sterile disks containing the antibiotic were placed on top. Plates were incubated at 37°C for 24 h and inhibition zones were measured using ImageJ. ZOI_resistant_ > ZOI_WT_ indicates CS and ZOI_resistant_ < ZOI_WT_ indicates CR. Such a profile would serve as a preliminary indicator for CR-CS. The extent of CS was expressed in terms of MIC_WT_: MIC_resistant_ strain.

### Hierarchical clustering of resistant isolates

The diameter of the zone of inhibitions was considered a quantitative trait and hierarchical clustering was performed with average linkage and Euclidian distance assessment. Heatmap and cladograms were generated to illustrate the clustering.

### Determining copy number variation by qPCR

QPCR has been implemented in determining gene copy number in *Acinetobacter* spp. [29]. Comparative copy number analysis was performed by qPCR with primers selective for *pmr*A and *pmr*B genes with an amplicon size of 150-180 bp with oligonucleotides enlisted in Table S-1. SYBR green-based qPCR was performed with 1 ng of genomic DNA preparation. Copy number change was scored based on ΔCt against gapdh gene amplification from the same preparation.

### Sanger sequencing

Sanger sequencing was performed for amplicons obtained from PCR targeting selective genes. PCR amplicons were gel-purified prior to sequencing. The Primer sequences are mentioned separately (Table S1).

### Testing the presence of mcr gene

For detecting the presence of mcr genes, primers were designed for the various *mcr* genes (mcr1-10), and PCR amplifications were performed from the genomic DNA isolated from the evolved and resistant lines using primers enlisted in Table S1.

### Whole-genome sequencing

Resistant mutant displaying significant CR/ CS was ranked according to the extent and number of antibiotics against altered susceptibility. 3 clonal populations obtained from independently evolved lines have been sent for whole genome sequencing. For genome sequencing, total DNAs were prepared using the QIAamp DNA Purification Kit. NexteraXT libraries were prepared and sequenced on an Illumina Novaseq 6000. Paired sequence reads will be mapped using BWA-MEM [30] to the Ab19606 genome. The variants were called from the aligned bam file using FreeBays available in BV-BRC [31].

### Biofilm assay

Biofilm assay was performed as described earlier [32]. Briefly, bacteria were inoculated from log phase cultures grown in Luria-Bertani broth with glucose at 37 °C till 0.6 OD_600_ was reached. The cultures were then distributed into the wells of 96-well polystyrene and will be incubated at 30 °C for 24 h discarding the media carefully, the wells were washed thrice with sterile distilled water. Biofilm formation at the liquid-substratum interface was quantified by measuring OD_570 nm_ for stain (crystal violate) retention by the biofilm. For each set of experiments an uninoculated well and a well, freshly filled with overnight culture, were used as negative control and a non-biofilm negative control system respectively.

To examine biofilm formation silicone surface associated with medical tubing, a catheter biofilm formation assay was performed as described elsewhere [15] and biofilm formation was observed by crystal violate retention.

### Viability in biofilm cells

Biofilms of Ab were allowed to develop in a liquid-polystyrene substratum interface for 24 h. The impact of antibiotic-pigment combinations on the viability of biofilm cells was estimated for the preformed biofilms by resazurin-reduction as described elsewhere [33]. Briefly, following removal of the exhausted medium, the preformed biofilms were washed twice with saline and were incubated in 200 µl of MHB with 25 µl of resazurin (200 µg/ ml) for 60 mins. The extent of resazurin reduction was monitored by estimating OD_570 nm_. CFU/ ml for the biofilms was enumerated by suspending the biofilm, dilution, and further plating.

### Autoaggregation assay

After growing bacteria in LB broth for 24 h at 37°C, cells were centrifuged and suspended in phosphate-buffered saline (PBS) to 0·5 OD units at 600 nm. 2 ml bacterial suspension was placed in each tube, centrifuged and then re-suspended in PBS. After 2-3 h exposure at 37°C, 1 ml of the upper layer of suspension was transferred to another tube and the OD_595 nm_ was measured. Aggregation was expressed as 1−(O.D_upper suspension_/O.D_total bacterial suspension_)×100.

### Hydrophobicity of the cell surface

A 1.2 ml volume of cell suspension in phosphate/urea/Mg (PUM) buffer, pH 7.5 (22.2 g K_2_HPO_4_. 3H_2_O, 7.26 g KH_2_PO_4_, 1.8 g urea, 0.2 g MgSO_4_. 6H_2_O, 1 l deionized water), was mixed with 200 ml of n-hexadecane with vigorous stirring for 2 min. After phase separation at room temperature for 15 min, the lower aqueous phase was removed and its OD_580nm_ was measured.

### Surface motility

Surface-assisted motility was visualized according to Mayer et al., 2018 [34]. One microliter from overnight cultures at 0.3 optical density (OD_600 nm_) was seeded in the center of the plates. Subsequently, plates were incubated at 37°C in the dark. Surface-associated motility trail was observed after 48 h.

### EPS extraction and assay

Bacterial biomass was carefully harvested from the biofilms developed in a solid-liquid interface and suspended in saline. Following dispersion by vigorous stirring and clarification by high-speed centrifugation the supernatant was filtered through cellulose acetate membranes to obtain a cell-free EPS solution.

Relative quantitation of eDNA was performed after extraction using a Soil DNA purification kit (HiMedia) by PCR amplification of *gapdh* and *16SrRNA*. The total carbohydrate content was estimated by the anthrone method as described earlier [35]. Briefly, 80 µl of the supernatant was mixed with 160 µl of anthrone reagent (0.125% Anthrone in 94.5 % H_2_SO_4_) and were incubated in the water bath at 100° C for 14 minutes and then cooled at 4°C. The OD_630 nm_ was measured using a microplate reader iMark microplate reader (Biorad).

### Outermembrane integrity

The integrity of the outer membrane was examined by N-phenyl-1-naphthylamine (NPN) - permeability assay according to Abdullah et al. [36]. Briefly, 10^6^ log phase bacteria were resuspended in 10 mM sodium phosphate buffer to achieve a final OD_600 nm_ 0.1 exposed to 0.2mM NPN in of and Em_420nm_ was measured (Ex at 350 nm).

### Statistical analysis

Statistical analysis was performed for the experiments with Students pairwise t-test. At least three technical replicates were tested for each biological replicate.

## Result

### Experimental evolution of colistin-resistant lines of A. baumannii19606

A single clonal population of Ab19606 was isolated by dilution plating and the population was used as a *source clonal population* for step-wise selection against increasing doses of COL and CIP (Fig. S1). MIC values of 0.625 µg/ ml and 1 µg/ ml for COL and CIP respectively against Ab were determined by microdilution (Table 1). Following each level of selection strain confirmation by Ab gene-specific primers (*gapdh* and/or 16SrRNA) was performed (Fig. S1). After final selection, the PCR amplicons were analyzed by Sanger sequencing followed by BLASTN against Ab19606 genome version AP022836.1. The time-line evolutionary selection trail, for the antibiotics was determined by measuring the time required to attain late log-phase (OD_600 nm_ = 0.8). As depicted in Fig. S2A, for CIP the selection pressure worked at its highest at the initial stages for selection (1X and 2X MIC) while it depleted in the following steps (Fig. S2A). For COL, the population adopted at the lowest rate at 1X MIC, though it remained slow-growing throughout the next levels of selection (Fig. 1A).

**Fig. 1.**
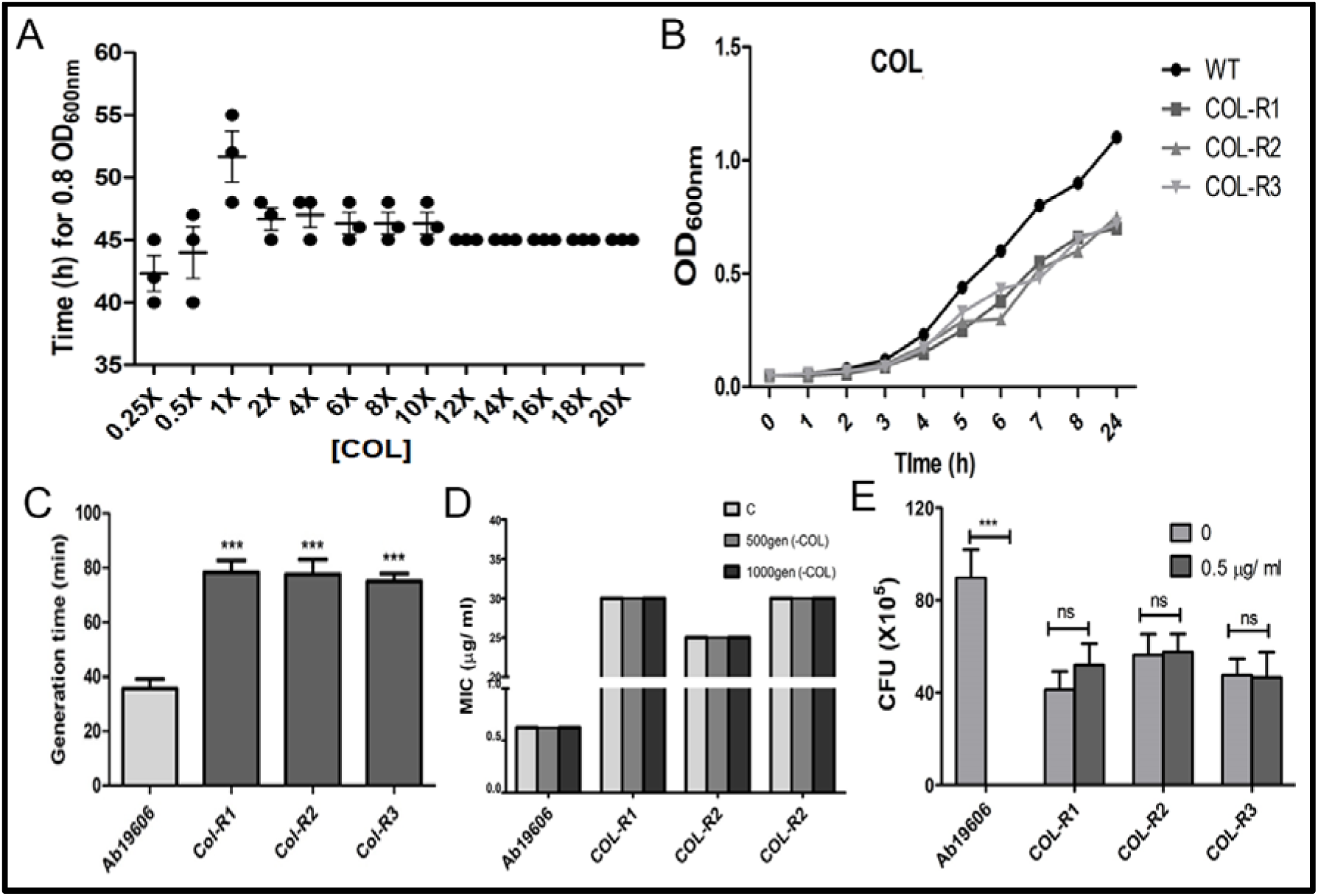
Experimental evolution of antibiotic-resistant lines and fitness of the evolved lines. Antibiotic-resistant lines were evolved by step-wise selection of a source clonal population against gradually increasing doses of antibiotic starting from 0.25XMIC (0.156 µg/ ml) for COL to a maximal selection at 20X MIC (12.5 µg/ ml) (A). Selection dynamics (trail) at each stage of variation is projected in the time needed to reach the late-log phase (OD600nm = 0.8) at each step of selection. Data represents mean ±SEM for three independent lines (n=3 for each). (B) Growth kinetics for the Ab source, AbCOL-R1, R2, and R3 clonal populations selected against COL were performed by monitoring OD600 nm for 24 h while growing in the absence of the antibiotic. Data represents mean ±SEM for three independent lines (n=3 for each). (C) Generation time for exponential growth for Ab source, AbCOL-R1, R2, and R3 clonal populations was determined by estimating the time needed for doubling in CFU/ ml value between 4h -7h post inoculation (log-phase). Data represents mean ±SEM for three independent lines (n=3 for each). ***p<0.001, two tailed paired Student’s t-test. (D) The stability of the resistance phenotype for AbCOL-R1, R2, and R3 was assessed by determination of MIC after nonselective growth for ∼500 generations and ∼1000 generations by serial subculture and comparing the value with MIC determined for populations (control, C) cultured with 0.5XMIC of COL for respective lines. A parallel set was replicated for the Ab source clonal population (Ab19606) cultured in the absence of COL. © COL-dependence of AbCOL-R1, R2, and R3 was examined by enumerating changes in CFU/ ml for AbCOL-R1, R2, and R3 when plated on MHA with or without 0.5 µg/ ml of COL. The source clonal population (Ab19606) was verified for susceptibility to COL under a similar set-up (n=3 for each). Ns, not significant, ***p<0.001, two-tailed paired student t-test.

**Table 1.** MIC of reference strain against the test. MIC (µg/ ml) for the reference *A. baumannii* 19606 and experimentally evolved colistin, ciprofloxacin, and vancomycin-resistant lines were determined by microdilution assay.

|  | **Ab19**<br>**606** | **COL-**<br>**R1** | **COL-**<br>**R2** | **COL-**<br>**R3** | **CIP-**<br>**R1** | **CIP-**<br>**R2** | **CIP-**<br>**R3** | **VNC-**<br>**R1** | **VNC-**<br>**R2** | **VNC-**<br>**R3** |
| --- | --- | --- | --- | --- | --- | --- | --- | --- | --- | --- |
| Colistin | 0.625 | 25 | 25 | 25 |  |  |  | 25 | 12.5 | 25 |
| Gentamycin | 1.5 | 0.8 | 0.8 | 0.4 |  |  |  |  |  |  |
| Vancomycin | 50 | 3.125 | 3.125 | 3.125 |  |  |  | >500 | >500 | >500 |
| Ciprofloxacin | 0.5 | 0.025 | 0.025 | 0.025 | 6 | 6 | 6 |  |  |  |
| Aztreonam | 50 | 25 | 25 | 25 |  |  |  |  |  |  |

### Growth, colistin dependence, and fitness of the evolved colistin-resistant lines

Growth of each evolved line in the absence of selection pressure was recorded by measuring OD_600 nm_ at specific time intervals. As represented in Fig. S2B, clonal populations derived from the lines evolved against CIP demonstrated a growth profile similar to Ab19606 with slopes 0.166, 0.193, and 0.184 against 0.167 for Ab19606 source clonal population for exponential growth respectively. The three evolved COL-R lines displayed a considerably reduced growth rate compared to the Ab19606 line (Fig. 1B), with mean slopes of 0.119, 0.1, and 0.109 against a mean slope of 0.167 for exponential growth. Generation time for three clonal populations for each of the AbCOL-R lines was determined by estimating CFU at a 3h time interval with an exponential phase of growth for each. As depicted in Fig. 1C, 2.22±0.22, 2.19±0.21, 2.13±0.24 –fold increase in generation time was estimated for AbCOL-R1, AbCOL-R2, and AbCOL-R3 clonal population against the generation time of 35.66±4.92 mins for Ab19606 clonal population (Fig. 1C) indicating significant (P<0.001, n=3) compromise of fitness associated with resistance evolution. To examine the stability of the resistant phenotype, three independent clonal populations from each of the AbCOL-R lines were grown without COL. Following growth of 500 and 1000 generations, the resistance phenotype was verified by determining MIC against COL, and no apparent alteration for MIC was observed for the resistant bacteria (Fig. 1D). COL-dependent growth has been reported for several extremely COL-resistant isolates of *A. baumannii* [37]. When AbCOL-R1, AbCOL-R2, and AbCOL-R3 clonal populations were grown on MHA plates in the absence of COL and the presence of COL (0.5 µg/ ml), no significant difference in CFU was noted for the two conditions (Fig. 1E), suggesting that the extremely resistant evolved strains were not dependent on COL for growth.

### Colistin-resistant mutants demonstrated hyperbiofilm formation

AbCOL-R1, R2, and R3 grew slowly compared to the source (Fig. 1B) and the evolved strains are profound biofilm formers. As determined by static biofilm formation assay on polystyrene substratum, clonal populations from AbCOL-R1, R2, and R3 forms profound biofilms compared to the source (4.77±0.24, 4.90±0.72, 4.93±0.49-fold respectively, p<0.001, Fig. 2A). The AbCIP-R1, AbCIP-R2, and AbCIP-R3 clonal population demonstrated biofilm forming potential similar to the Ab19606 source clonal population with 1.16±0.06, 1.09±0.36, and 1.16±0.30-fold change in crystal violate retention (Fig. S3). The static biofilm forming potential of the AbCOL-R1 and AbCOL-R2 strains were further validated in an *in vitro* catheter surface biofilm formation model where the evolved lined showed markedly enhanced adherence and film development on catheter wall compared to Ab19606 (Fig. 2B). To evaluate bacterial autoaggregation, sedimentation assays were performed with late log-phase cultures of Ab, AbCOL-R1, R2, and R3 by measuring the OD_600nm_ value of the bacterial culture supernatant after 1 h of settling period. Compared to Ab19606, AbCOL-R1, and R2 demonstrated 55.26±2.57% and 28.57±5.69% aggregation. No significant difference was noted for AbCOL-R3 compared to the source clonal population (Fig. 2C). To characterize biofilm development in the evolved lines further, EPS was extracted from the submerged biofilms formed by clonal populations, and total polysaccharide and eDNA were estimated. As resented in Fig. 2D, 4.18±0.52, 5.03±0.78, and 5.43±0.94 fold production of EPS polysaccharide was determined for biofilms formed by AbCOL-R1, AbCOL-R2, and AbCOL-R3 respectively. Indirect quantification of eDNA from the EPS preparations by PCR for *gapdh* genes indicated 1.88±0.12, 2.02±0.14, and 1.85±0.13-fold enrichment of eDNA in the EPS preparations derived from biofilms formed by AbCOL-R1, AbCOL-R2, and AbCOL-R3 respectively (Fig. 2E). Cell surface hydrophobicity has been identified as a major determinant of biofilm development in *A. baumannii* [38]. Comparative analysis of cell surface hydrophobicity indicates that the surface of clonal populations derived from AbCOL-R1, and R2 are 2.13±0.02 and 1.49±0.04-fold more hydrophobic respectively compared to the source Ab19606 clonal population (Fig. 2F). At the same time, no significant difference could be detected for AbCOL-R3.

**Fig. 2.**
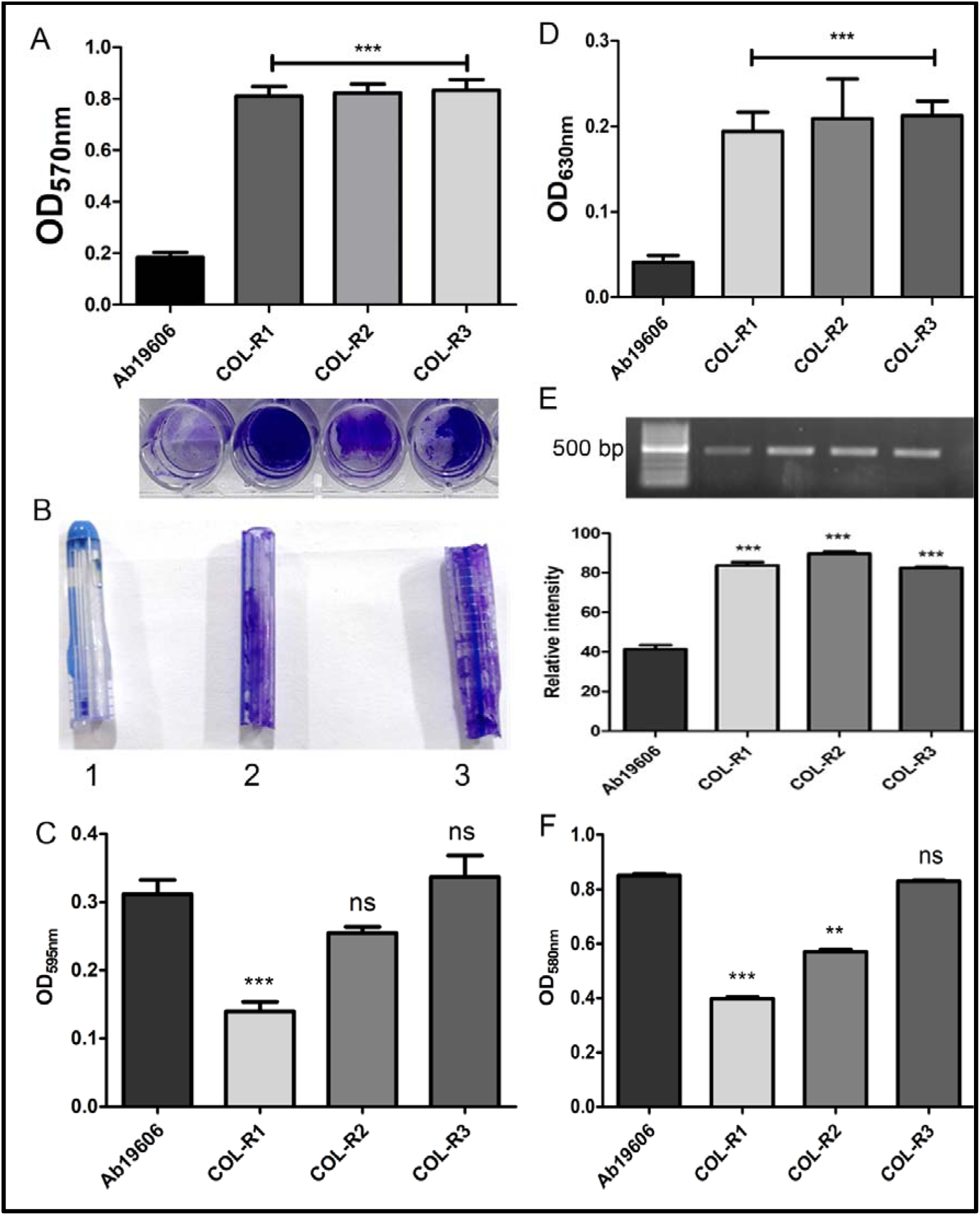
Biofilm development by COL-R mutants. (A) Static biofilm formation on polystyrene substratum was envisioned by crystal violate retention for each of the Ab source populations (Ab19606), AbCOL-R1, R2, and R3. Results represent mean±SEM for three independent experiments. ***p<0.001, two-tailed paired Student t-test. In the set: representative data from one replicate of the experiment demonstrating biofilm development. (B) Log-phase cultures of Ab19606, AbCOL-R1, and AbCOL-R2 were allowed to form biofilm on the catheter interior surface. The extent of bacterial biofilm formation was envisioned by crystal violate staining. (C) Log-phase Ab source population (Ab19606), AbCOL-R1, R2, and R3 bacterial population were allowed to form an aggregate in a sedimentation buffer for 2h. Autoaggregation was estimated by measuring OD600nm of the top layer of the suspension. Results represent mean±SEM for three independent experiments. Ns, not significant, ***p<0.001, two-tailed unpaired Student’s t-test. (D) Late log phase cultures of Ab19606, AbCOL-R1, R2, and R3 were allowed for biofilm at the liquid-polystyrene substratum interface. EPS was extracted from the matured biofilms and total EPS polysaccharide was estimated by the anthrone acid method (n=3, for each). Results represent mean±SEM for three independent experiments. ***p<0.001, two-tailed unpaired Student’s t-test. © eDNA was indirectly quantitated by performing PCR with gapdh gene-specific primers with an amplicon size of 500 bp, subsequent agarose gel electrophoresis, and densitometric analysis. Data represent mean±SEM for three independent experiments. ***p<0.001, two-tailed unpaired Student’s t-test. (F) Cell surface hydrophobicity was compared for each treatment after allowing log-phase bacterial cells of the source population (Ab19606), AbCOL-R1, R2, and R3 suspended in PUM buffer to accumulate into the n-hexadecane phase. Results represent mean±SEM for three independent experiments. Ns, not significant, ***p<0.001, **p<0.01, two tailed unpaired Student’s t-test.

To determine the viability of biofilm cells, conversion of resazurin to resorufin was tracked for biofilms formed by AbCOL-R1, AbCOL-R2, and AbCOL-R3, and 2.84±0.53, 2.74±0.11, 3.05±0.42-fold (P<0.001, n=3 for each) greater conversion was noted compared to biofilm cells of the source clonal population (Fig. 3A). Biofilm formation in *A. baumannii* is linked with regulation of surface motility [39]. The surface motility of the evolved lines was examined on motility agar (0.5%) plates following a spot inoculation and the maximal distance of the front in any direction was enumerated after 30h. As depicted in Fig. 3B, AbCOL-R1, AbCOL-R2, and AbCOL-R3 were substantially compromised in surface motility with ∼85.64%, ∼90.83%, and ∼88.80% decline in distance traveled compared to Ab19606 (Fig. 3B). The results indicated enhanced biofilm development, along with retarded growth rate, as a significant trade-off in experimental evolution of COL-resistance. To further explore the trade-off, the expression of three genes, ascribed to biofilm development and maturation in Ab, namely *bap*, *ompa*, *csu*E were analyzed from AbCOL-R1. As presented in Fig. 3C, the expression of *ompa* remained unaltered compared to the source clonal population, while the expression *bap* was determined to be down-regulated by 2.1±0.39 –fold. Despite being a major facilitator for biofilm development, expression of *csuE* was markedly down-regulated by 8.66±0.14 –fold in the hyperbiofilm-forming AbCOL-R1 clonal populations (Fig. 3C).

**Fig. 3.**
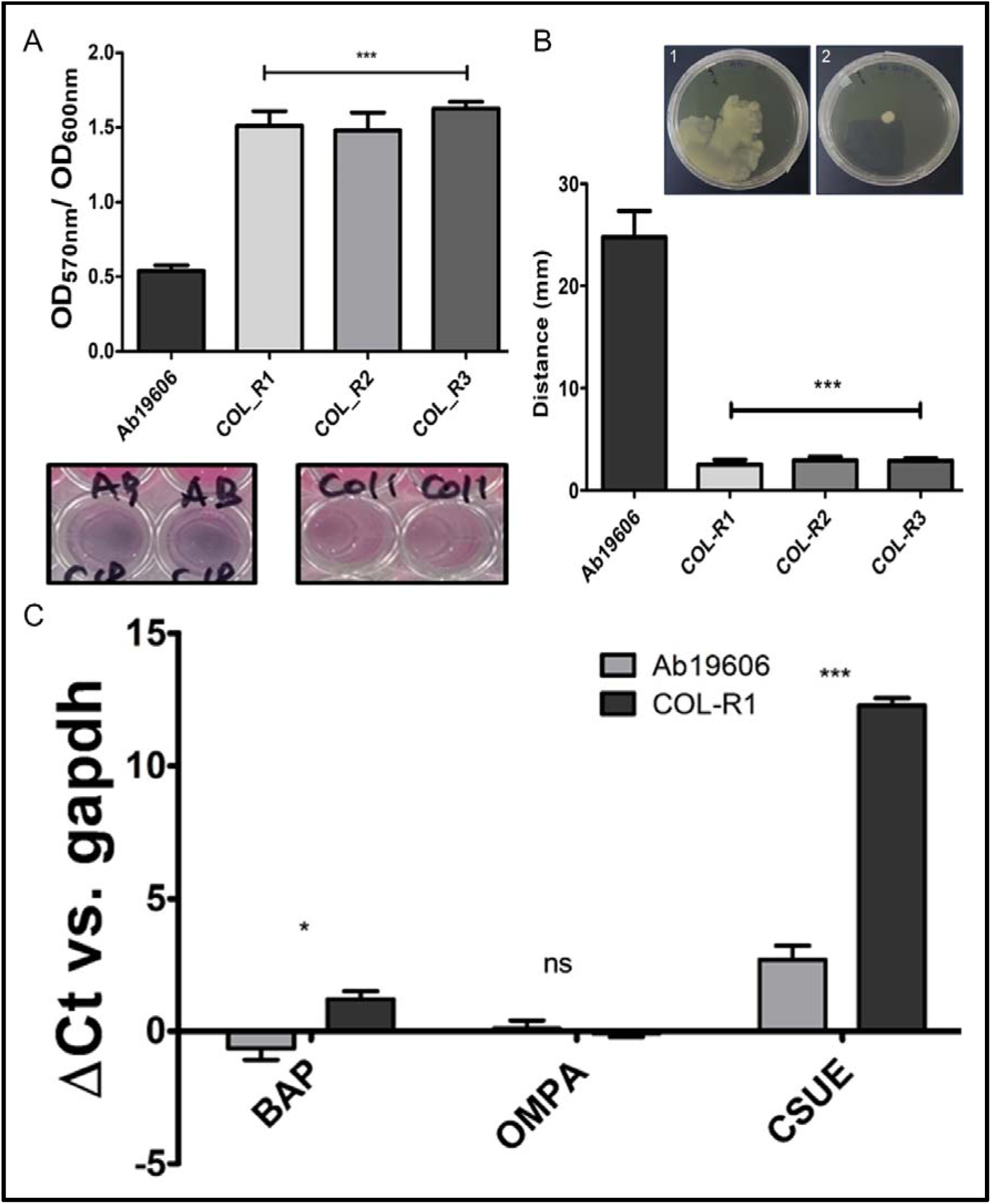
Biofilm-associated attributes for COL-resistant mutants. (A) The viability of biofilm cells for the Ab source population (Ab19606), AbCOL-R1, R2, and R3 bacterial population was estimated after allowing late log-phase Ab source population (Ab19606), AbCOL-R1, R2, and R3 bacterial population to form biofilms and enumerating metabolic activity of the biofilms by resazurine assay. Results represent mean±SEM for three independent experiments. ***p<0.001, two-tailed unpaired Student’s t-test. (B) Surface motility of Ab19606, AbCOL-R1, R2, and R3 was envisioned by spot inoculation on 0.5% motility agar from log phase cultures of the lines and tracking motility for 48 h of the distance of the progressive-front for each (n=3). Results represent mean±SEM for three independent experiments. ***p<0.001, two tailed paired Student’s t-test. (C) Analysis of expression for bap, ompa, and csuE genes in late-log phase Ab19606, AbCOL-R1, R2, and R3 clonal population was performed by qPCR. Expression with respect to gapdh expression is expressed as ΔCt. Results represent mean±SEM for three independent experiments. Ns, not significant, ***p<0.001, two-tailed unpaired Student’s t-test.

### Colistin-resistant mutants are impaired of outer-membrane integrity

As a primary characterization at the molecular level known genetic markers for COL-resistance line presence of *mcr* genes, and copy number variation (CNV) of single nucleotide variation (SNV) in *pmr*CAB and *lpx*ABC loci were analyzed by PCR amplification, qPCR, and by whole genome sequencing. The outer membrane synthesis linked *pmr* and *lpx* genes have been associated with COL resistance [5,40] along with *mcr* genes encoding an array of phosphoethanolamine transferase [22]. Various *mcr* genes (*mcr*1-10) were amplified with specific primers with amplicon sizes ranging from 322 to 1644 bp. For *mcr*1 and *mcr*2 positive control isolates were available in the laboratory. For all, amplification was performed for genomic DNA isolated from AbCOLR-1, AbCOLR-2, and AbCOLR-3. As depicted in Fig. 4A, none of the evolved strains harbored any of the *mcr* genes (Fig. 4A). The copy number for *pmr*A and *pmr*B genes were enumerated from the isolated genomic DNAs of the resistant strains by qPCR. As demonstrated in Fig. 4B, the ΔCt value against *gapdh* for *pmr*A were 1.29±0.12, 1.21±0.18, 1.10±0.09, and 1.16±0.16 for Ab19606, AbCOLR-1, AbCOLR-2, and AbCOLR-3 respectively. Similarly, for *pmr*B, the ΔCt value against *gapdh* were 1.40±0.07, 1.28±0.12, 1.14±0.10, and 1.24±0.066 for Ab19606, AbCOLR-1, AbCOLR-2, and AbCOLR-3 respectively, indicating no apparent change in copy number (Fig. 4B). To identify genomic variations associated with experimental evolution of COL resistance, WGS was performed with the DNA isolated from single clonal population from AbCOL-R1, AbCOL-R2, and AbCOL-R3 along with the source clonal population of Ab19606. As enlisted in Table-XX a number of variants could be called in the mutant genomes which included both InDels and nonsynonymous SNPs. Among the high-confidence (score>2500) nonsynonymous SNPs identified, the lpxA2 gene has earlier been ascribed to COL resistance. The InDel identified resulted in a duplication of proline (P190), rendering possible disruption of the structure of the catalytic core (Table 2). Alongside, a G87C mutation was identified hvrA, and a putative DNA binding protein, R1136C mutation was predicted in rpoC gene encoding DNA-directed RNA polymerase β’ subunit. A S34L mutation was identified as a common variation in sigma-fimbriae tip adhesion protein. In addition, in AbCOL-R1 and AbCOL-R2, L1F substitution was enlisted and A220T substitution was identified in a transcriptional regulator of the AraC family in AbCOL-R3 (Table 2). The integrity of the outer membrane was assessed by NPN assay with log-phase culture of Ab source, AbCOL-R1, R2, and R3 clonal population. As depicted in Fig. 4C, 1.79±0.06, 1.74±0.13, and 2.28±0.17 –fold increase in emission (420 nm) were recorded for AbCOL-R1, AbCOL-R2, and AbCOL-R3 respectively compared to the Ab source clone population (Fig. 4C). The result indicated impairment of outer membrane in the resistant mutants.

**Fig. 4.**
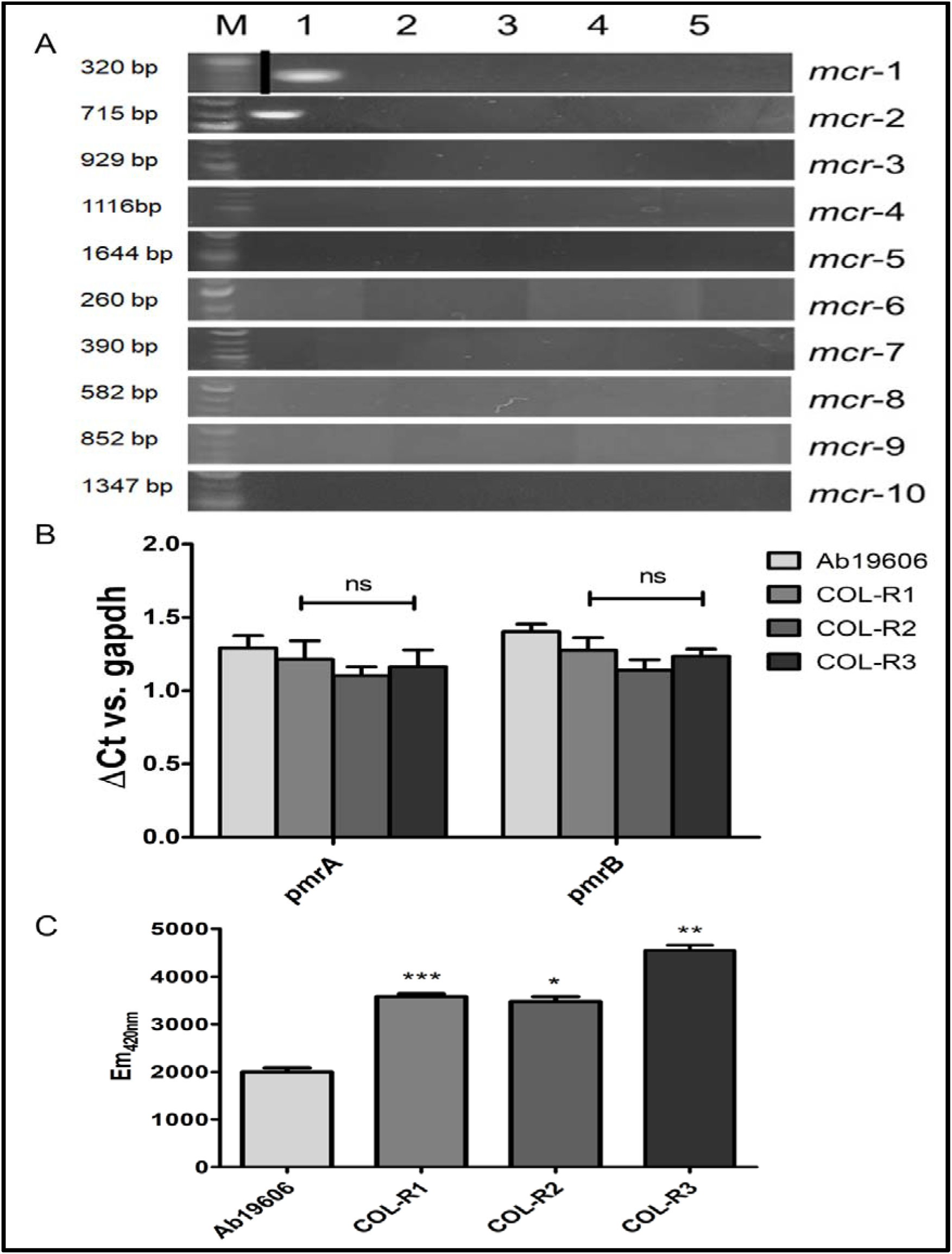
Analysis of COL-resistance markers and outer membrane integrity. (A) The presence of extrachromosomal transmitted COL-resistance marker mcr genes was profiled in Ab19606 (2), AbCOL-R1 (3), AbCOL-R2 (4), and AbCOL-R3 (5) by mcr-specific primers (mcr1 to 10). Specific amplicon sizes for each of the mcr are mentioned and detected with respect to the 1 kb DNA ladder (M). Positive controls for known DNA containing specific mcrs were available for mcr1 and mcr2 only (1). (B) Copy number variations for pmrA and pmrB were examined by qPCR with specific markers for the genes and genomic DNA isolated from the Ab19606, AbCOL-R1, R2, and R3. CNV was expressed in terms of ΔCt against gapdh amplification. Results represent mean±SEM for three independent experiments. Ns, not significant, two-tailed unpaired Student’s t-test. (C) The integrity of the outer membrane for Ab19606, AbCOL-R1, R2, and R3 was examined by NPN permeability assay. Log phase bacteria were exposed to 0.2mM of N-phenyl-1-naphthylamine (NPN) and Em420nm was measured (Ex at 350 nm). Results represent mean±SEM for three independent experiments. ***p<0.001, **p<0.01, *p<0.05, two tailed unpaired Student’s t-test.

**Table 2.**
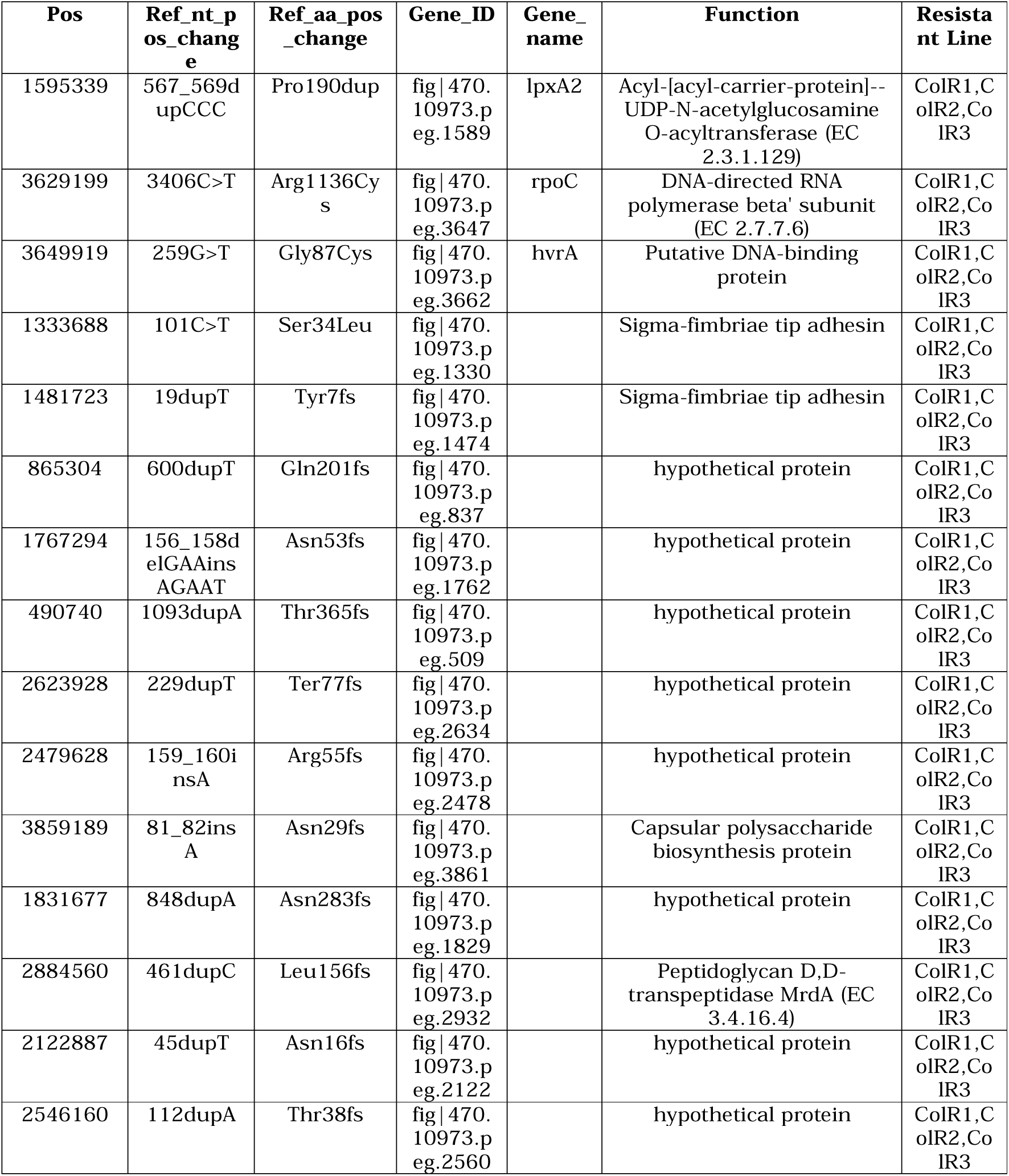
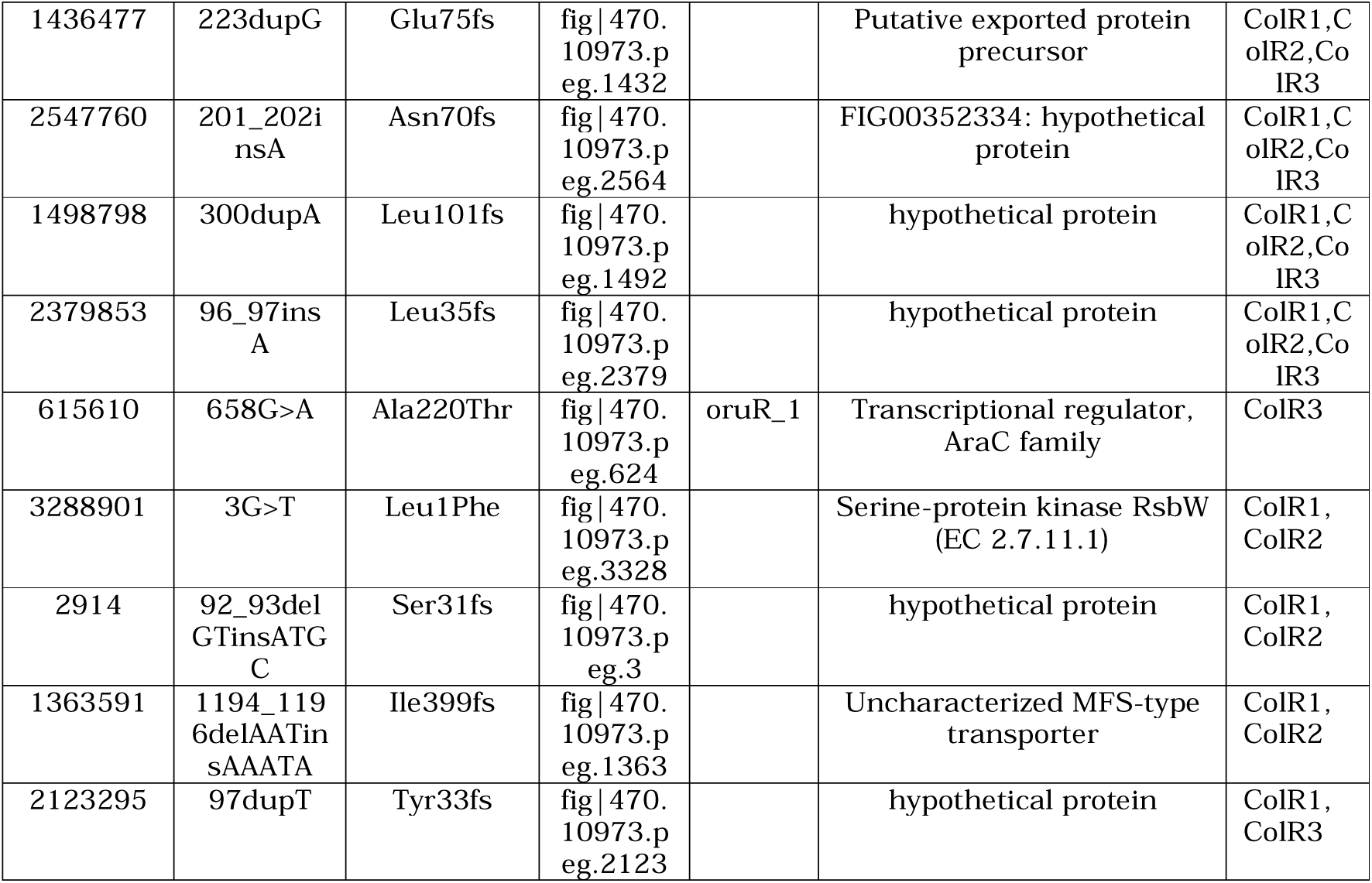
Intragenic variation identified in colistin-resistant mutants. Whole genome sequencing was performed from clones derived from three colistin-resistant lines (Col-R1, R2, and R3) and the source clonal population. Reads were aligned with *A. baumannii* 19606 reference genome and single nucleotide variants (SNV) were called. For the unique SNVs identified in COL-R mutants, positions in chromosomes, changes in nucleotide, and changes in amino acid, gene ID, gene names, and functions are enlisted. The clones in which the variations were identified are also mentioned.

| Pos | Ref_nt_pos_change | Ref_aa_pos_change | Gene_ID | Gene_name | Function | Resistant Line |
| --- | --- | --- | --- | --- | --- | --- |
| 1595339 | 567_569dupCCC | Pro190dup | fig\|470.10973.p eg.1589 | lpxA2 | Acyl-[acyl-carrier-protein]--UDP-N-acetylglucosamine O-acyltransferase (EC 2.3.1.129) | ColR1, ColR2, ColR3 |
| 3629199 | 3406C>T | Arg1136Cys | fig\|470.10973.p eg.3647 | rpoC | DNA-directed RNA polymerase beta' subunit (EC 2.7.7.6) | ColR1, ColR2, ColR3 |
| 3649919 | 259G>T | Gly87Cys | fig\|470.10973.p eg.3662 | hvrA | Putative DNA-binding protein | ColR1, ColR2, ColR3 |
| 1333688 | 101C>T | Ser34Leu | fig\|470.10973.p eg.1330 |  | Sigma-fimbriae tip adhesin | ColR1, ColR2, ColR3 |
| 1481723 | 19dupT | Tyr7fs | fig\|470.10973.p eg.1474 |  | Sigma-fimbriae tip adhesin | ColR1, ColR2, ColR3 |
| 865304 | 600dupT | Gln201fs | fig\|470.10973.p eg.837 |  | hypothetical protein | ColR1, ColR2, ColR3 |
| 1767294 | 156_158delGAAinsAGAAT | Asn53fs | fig\|470.10973.p eg.1762 |  | hypothetical protein | ColR1, ColR2, ColR3 |
| 490740 | 1093dupA | Thr365fs | fig\|470.10973.p eg.509 |  | hypothetical protein | ColR1, ColR2, ColR3 |
| 2623928 | 229dupT | Ter77fs | fig\|470.10973.p eg.2634 |  | hypothetical protein | ColR1, ColR2, ColR3 |
| 2479628 | 159_160insA | Arg55fs | fig\|470.10973.p eg.2478 |  | hypothetical protein | ColR1, ColR2, ColR3 |
| 3859189 | 81_82insA | Asn29fs | fig\|470.10973.p eg.3861 |  | Capsular polysaccharide biosynthesis protein | ColR1, ColR2, ColR3 |
| 1831677 | 848dupA | Asn283fs | fig\|470.10973.p eg.1829 |  | hypothetical protein | ColR1, ColR2, ColR3 |
| 2884560 | 461dupC | Leu156fs | fig\|470.10973.p eg.2932 |  | Peptidoglycan D,D-transpeptidase MrdA (EC 3.4.16.4) | ColR1, ColR2, ColR3 |
| 2122887 | 45dupT | Asn16fs | fig\|470.10973.p eg.2122 |  | hypothetical protein | ColR1, ColR2, ColR3 |
| 2546160 | 112dupA | Thr38fs | fig\|470.10973.p eg.2560 |  | hypothetical protein | ColR1, ColR2, ColR3 |
| 1436477 | 223dupG | Glu75fs | fig\|470.10973.p<br>eg.1432 |  | Putative exported protein precursor | ColR1, ColR2, ColR3 |
| 2547760 | 201_202insA | Asn70fs | fig\|470.10973.p<br>eg.2564 |  | FIG00352334: hypothetical protein | ColR1, ColR2, ColR3 |
| 1498798 | 300dupA | Leu101fs | fig\|470.10973.p<br>eg.1492 |  | hypothetical protein | ColR1, ColR2, ColR3 |
| 2379853 | 96_97insA | Leu35fs | fig\|470.10973.p<br>eg.2379 |  | hypothetical protein | ColR1, ColR2, ColR3 |
| 615610 | 658G>A | Ala220Thr | fig\|470.10973.p<br>eg.624 | oruR_1 | Transcriptional regulator, AraC family | ColR3 |
| 3288901 | 3G>T | Leu1Phe | fig\|470.10973.p<br>eg.3328 |  | Serine-protein kinase RsbW (EC 2.7.11.1) | ColR1, ColR2 |
| 2914 | 92_93delGTinsATGC | Ser31fs | fig\|470.10973.p<br>eg.3 |  | hypothetical protein | ColR1, ColR2 |
| 1363591 | 1194_1196delAATinsAAATA | Ile399fs | fig\|470.10973.p<br>eg.1363 |  | Uncharacterized MFS-type transporter | ColR1, ColR2 |
| 2123295 | 97dupT | Tyr33fs | fig\|470.10973.p<br>eg.2123 |  | hypothetical protein | ColR1, ColR3 |

**Table 3.** MIC and FICI values for antibiotic combinations. MIC values for colistin (COL) and vancomycin (VNC) independently, COL in combination with VNC, and VNC in combination with COL were determined against *A. baumannii* 19606. A FICI value of ≤0.5 indicates synergy between the two antibiotics.

| **Antibiotic**<br>**and**<br>**combination** | **MIC**<br>**($\mu\text{g}/\text{m}$)** | **FICI** |
| --- | --- | --- |
| COL | 0.625 | 0.5 |
| COL(+VNC) | 0.156 |  |
| VNC | 50 |  |
| VNC(+COL) | 12.5 |  |

### Colistin resistance mutants demonstrated cross-resistance and collateral sensitivity against antibiotics CR

A comprehensive analysis of antibiotic responsiveness was performed for AbCOL-R1, AbCOL-R2, and Ab_COLR3 against 27 antibiotics representing all major groups of antibiotics. The lines demonstrated cross-resistance against and aztreonam while collateral sensitivity (CS) was observed against CIP, cefepime, chloramphenicol (CAM), fosfomycin, and vancomycin (VNC) compared to source Ab19606 (Fig. 5A). Intriguingly, *A. baumannii* is reportedly resistant to VNC [41] and that has been corroborated for Ab19606 (MIC 25-50 µg/ ml). The CS against VNC was validated by determining MIC, which showed ∼25-12.5-fold resensitization in AbCOL-R strains (Figs. 5B). The failure of the evolved COL-R strains to grow in Leeds *Acinetobacter* agar (Table-1) possibly can be attributed to the sensitivity to VNC [42]. To examine whether resistance emergence against VNC can demonstrate CS against COL, a VNC-resistant population was evolved by step-wise selection up to 500 µg/ ml. The AbVNC-R clonal populations did not demonstrate CS against CL, with no significant difference in MIC as determined by microdilution (0.6 to 1.2 µg/ ml vs. 0.3 to 0.6 µg/ ml, Fig. S4) and E-test (Fig. 5C). CS was also observed for CIP and fosfomycin, which was confirmed by MIC determination by microdilution demonstrating ∼20-fold sensitization against CIP (Table-1) and E-strip assay demonstrating significant sensitization (intrinsic resistant in Ab19606 vs. MIC of 2 µg/ ml in COL-R) against fosfomycin (Fig. S5). . Determination of MIC for gentamicin for which no significant CR or CS was determined by agar diffusion, indicated similar MIC (∼1.3 µg/ ml) for the clonal populations from evolved lines and the source clonal population of Ab19606 (Table-1). Though disc diffusion assay indicated modest CR against aztreonam for AbCOL-R mutants with slight increase in ZoI, the observation was not corroborated by MIC determination by microdilution (Table-1).

**Fig. 5.**
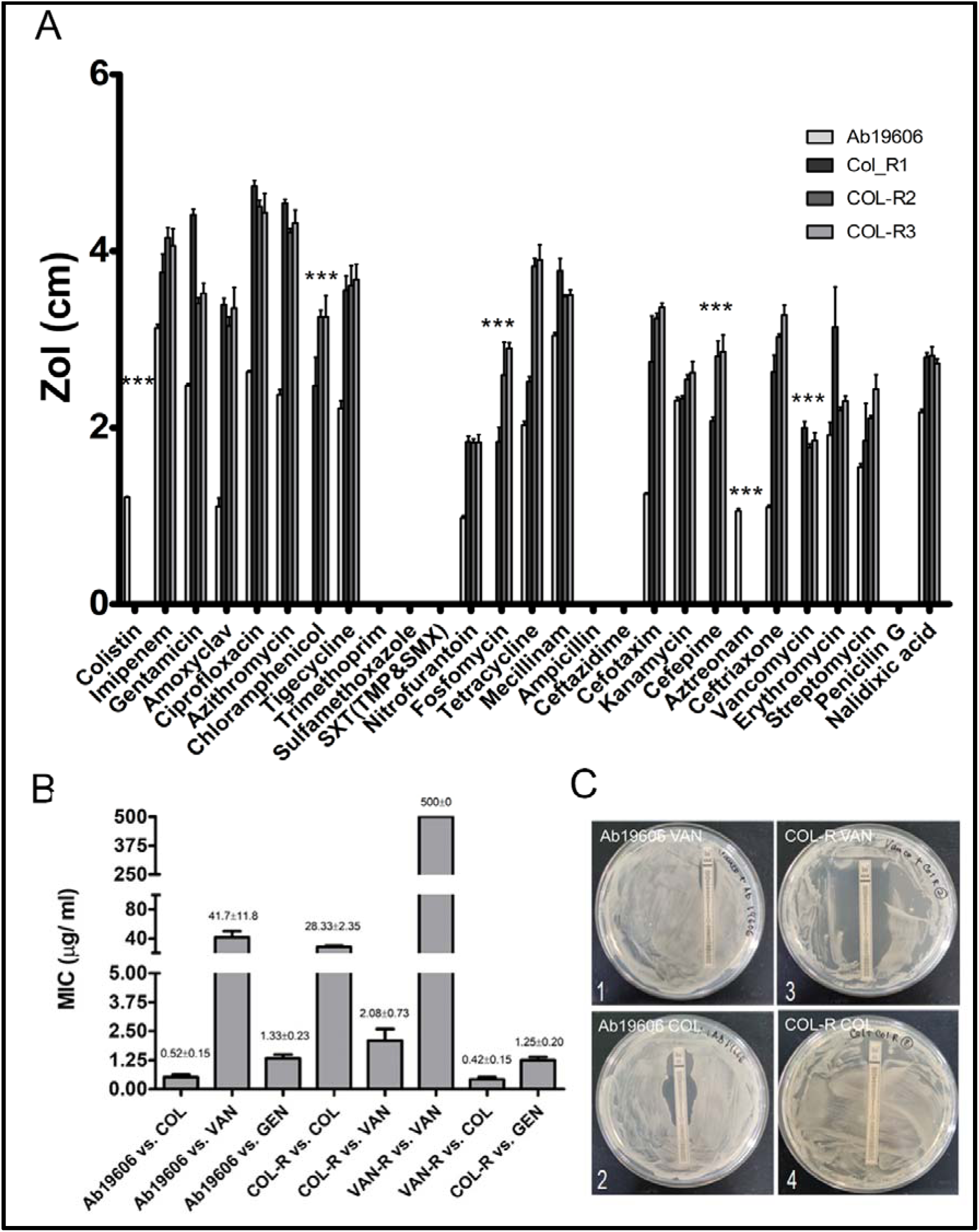
Comprehensive profiling for cross-resistance and collateral sensitivity against antibiotics for COL-resistant mutants. (A) The Kirby-Bauer disc diffusion-based CR-CS profiling was performed against 27 antibiotics for Ab19606 and AbCRP-R1, R2, and R3 lines. Data represent cumulative mean obtained from the three mutants±SEM for the three clones obtained from the lines. ***p<0.001, two-tailed paired Student t-test. (B) MIC against COL, VNC, and gentamicin (GEN) was determined for COL and VNC-resistant lines (COL-R and VNC-R) and Ab19606 cultured in parallel while generating resistant lines by microdilution. Results are represented as mean±SEM as observed from three independent experiments. (C) MIC against COL and VNC was also determined for An19606 and AbCOL-R1 by an E-test using MIC strips to confirm collateral sensitivity to VNC for AbCOL-R mutants. Representative images from three independent tests are included.

### Colistin-resistant cliniacal isolate demonstrated features similar to the experimentally evolved line

Three COL-resistant clinical isolates (DLPL20, DLPL37, and DLPL47) were availed and characterized for systemic antibiotic sensitivity trade-offs against 27 antibiotics comprising all major groups of antibiotics. Primary characterization against 12 antibiotics by VTEK2 reported DLPL20 to be resistant to pipericillin/ tazobactam, ceftazidime, meropenem, amikacin, gentamicin, levofloxacin, ciprofloxacin, and minocycline along with resistance against COL. DLPL37 demonstrated resistance against piperacillin/ tazobactam and COL (Table S2). DLPL47 demonstrated resistance only against COL (Table S2). Further characterization of antibiotic resistance by agar diffusion indicated that DLPL37 was resistant against ampicillin, penicillin G, and ceftazidime, and susceptibility against VNC, azithromycin, CIP, imipenem, and tigecycline (Fig. 6A). DLPL20 is a MDR strains resistant against majority of the test antibiotics. For DLPL47 susceptibility was noted against majority of the test antibiotic except mecillinam and ciprofloxacin (Fig. 6A). Among the isolates, DLPL37 showed biofilm formation and aggregation trade-offs similar to the experimentally evolved resistant lines (Fig.6B). The strain forms robust biofilms (∼4.1-fold, p<0.001) compared to Ab19606 (Fig. 6C).

**Fig. 6.**
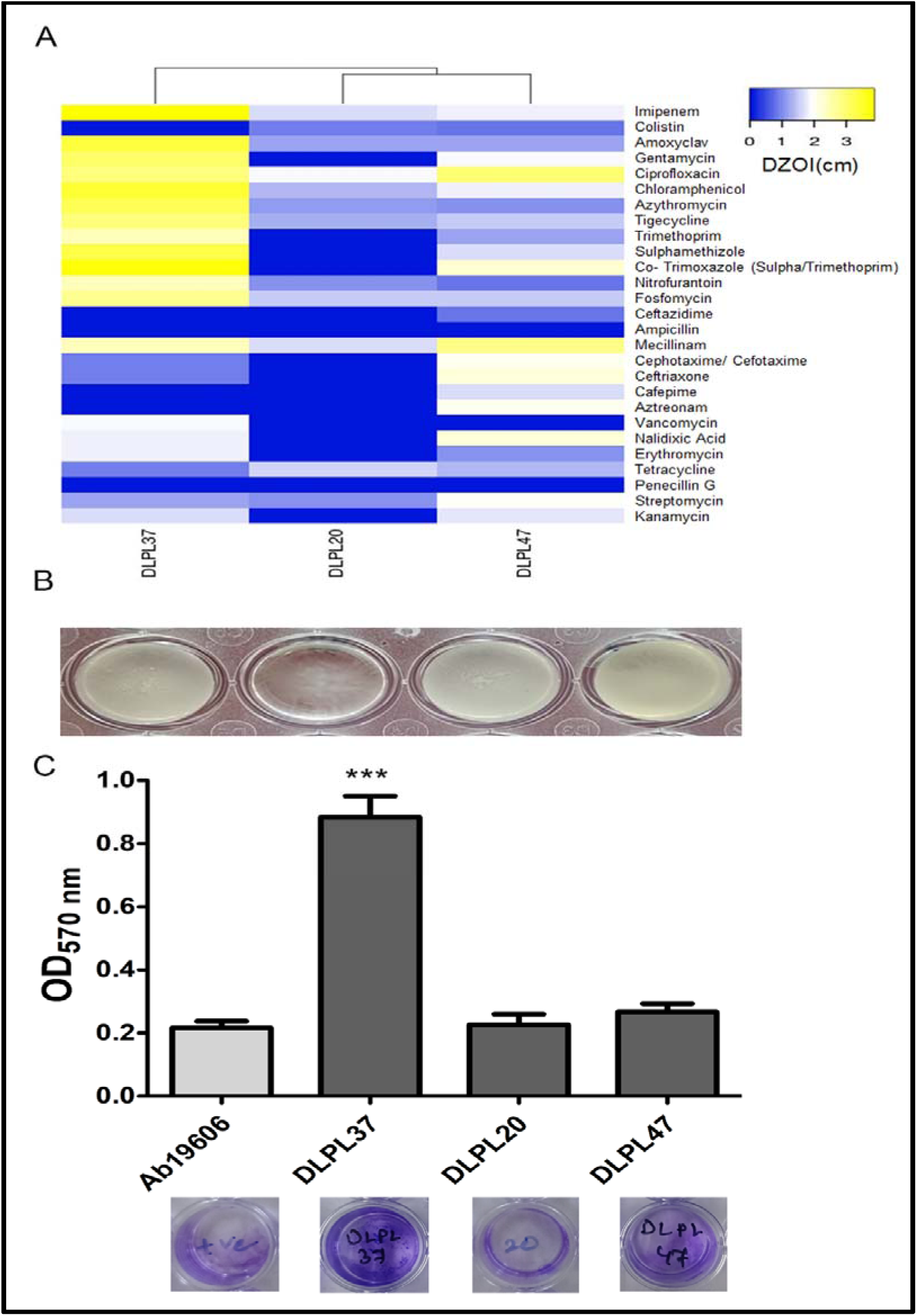
(A) Heat-map representation of the zone diameters (cm) as obtained from Kirby-Bauer disc diffusion based CR-CS profiling was performed against 27 antibiotics for DLPL20, DLPL37, and DLPL47. Data represent the cumulative mean obtained from the inhibition zone diameter from three experiments for the three resistant isolates. Hierarchical clustering was also performed for the isolates on the basis of the data obtained. (B) Late log-phase cultures of DLPL20, DLPL37, and DLPL47 were allowed to form biofilm. Visible aggregation and adherence were observed for DLPL37. (C) Static biofilm formation on polystyrene substratum was envisioned by crystal violate retention for each of the COL-resistant isolates. Results represent mean±SEM for three independent experiments (n=3). ***p<0.001, two-tailed paired Student t-test.

## Discussion

Owing to its tremendous genomic plasticity, Ab can promptly adapt to altered environmental conditions [43]. Such adaptations are commonly associated with phenotypic trade-offs influencing the physiology, metabolism, and fitness of bacteria in a broader sense [44]. Association of loss of LPS with COL resistance in *A. baumannii* has been described earlier with several mutations in lpxA, lpxB, and lpxC genes [12]. Alongside, variations in lipid A phosphoethanolamine (PEtN) transferase pmrC [5] and the two-component system associated with its expression of LPS, pmrA-B have been identified in COL-resistant mutants [45]. Plasmid-mediated polymyxin resistance (encoded by *mcr,* phosphoethanolamine transferase enzymes), particularly mcr-1 and mcr-4.3 has been revealed recently [21]. Here we aim to experimentally evolve COL-R-resistant mutants and a few clinical isolates to systematically profile cross-resistance and collateral sensitivity against antibiotics and other phenotypic trade-offs. The three experimentally evolved COL-R mutants were compromised in terms of LPS content. Of the previously reported COL-resistance-associated genes, variation of which is frequently attributed to loss or modification of LPS in Ab, in the three COL-R evolved mutants, only a single mutation in the lpxA2 gene could be identified. The mutation is an insertion resulting in the duplication of a proline (P190dup) residue. Whether the mutation impacts the function of lpxA2 warrants further experimentation. Among other common variations, the nonsynonymous substitution of the hvrA gene leading to G87C mutations warrants further exploration to identify whether such mutations are compensatory mutations or directly associated with COL resistance. No copy number variation for pmr genes could be observed in the mutants.

The impact of COL-R mutation on fitness and virulence attributes has been explored in various mutants and clinical isolates both in vitro and in vivo. In general, the term fitness is evaluated with comparative growth kinetic profiling and change in generation time. Mu et al reported variation in fitness cost among different COL-R mutants linked to genomic variation associated with COL resistance evolution [46]. Lower fitness in COL-R clinical isolates has been observed consistently [7,47]. The majority of the mutations identified in pmrA loci including E8D as associated with greater fitness cost. Tough P233S mutation in pmrB is not linked to fitness cost, other COL-R associated with the genomic variations inducted fitness cost [7]. Deletion, nonsynonymous, and frame-shift mutations associated with lpxA, C, and D are associated with fitness cost and virulence [48]. Corroborating the observation, the COL-R mutants evolved in this study by step-wise selection under submerged conditions demonstrated a significant fitness cost of ∼2-fold. However, whether the major mutation identified here in lpxA2 gene P190dup in terms of COL resistance or the Arg1136Cys mutation in DNA-directed RNA polymerase beta subunit (rpoC) is responsible for the cost, needs intricate exploration. Disruptive mutations of insertional elements IS*Aba1*, IS*Ajo2*, and ISAba13 leading to perturbed LPS and lipooligosaccharide synthesis render COL susceptible population to COL-dependent growth [37]. In the experimentally evolved COL-R mutants, no such insertions could be detected and the mutants did not display typical features of COL-dependent growth.

*Acinetobacter* sp. can form biofilm in various substratum at both solid-liquid and air-liquid interface [49]. Compared to other species, the intrinsic biofilm formation rate at the solid-liquid interface is around 3-times higher for Ab [50,51]. Antibiotic-resistant Ab isolates including CRABs have been reported to be prolific biofilm formers [50,52]. Moreover, higher expression levels for biofilm-associated genes like ompA, bap, csuE, epsA, blaPER−1, and bfmS expression have been observed in clinical isolates from diverse regions [53]. However, for COL-R strains, biofilm deficiency has been reported previously [47]. For both laboratory-induced and clinical isolates considerable depreciation in biofilm formation has been detected earlier and associated with LPS deficiency [26]. Contrary to earlier observations, in this study for AbCOL-R1-R3 robust biofilm formation was observed accompanied by significantly reduced surface motility. Moreover, severe depletion in the expression of *csuE* and significant depletion of *bap* expression was detected in the mutants. The results suggest that biofilm formation by the COL-R mutants is accomplished by some alternate mechanism involving the formation of a robust EPS enriched in polysaccharides and eDNA. Also, the mutation S34L mutation, identified in sigma-fimbriae tip adhesion protein, might be responsible for enhanced aggregation and adherence.

In clinical isolates of Ab, COL-resistance coupled with LPS-loss inflicted hypersensitivity to azithromycin, rifampicin, and VNC [54]. Among the laboratory-evolved strains of Ab17978, for several LPS-deficient strains 16arbouring a mutation in lpx genes, sensitivity was recorded against CIP, gentamicin, amikacin, minocycline, tigecycline, ampicillin, meropenem, and imipenem, while LPS modified strain demonstrated cross-resistance against most of the antibiotics [46]. For single strep selected LPS-deficient COL-R mutants hypersusceptibility was observed against VNC, rifampicin, and azithromycin, while against tigecycline and CIP the mutants were cross-resistant and minimal variation was observed for other β-lactam, carbapenems, amikacin, minocycline, and doxycycline [26]. For LPS-modified COL-R strains 16arbouring a mutation in pmrB, no significant difference was observed in antibiotic responsiveness [26]. Such reports clearly emphasized the heterogeneity among COL-R mutants in terms of antibiotic sensitivity trade-offs. In this study, we performed a comprehensive screening for CR-CS profiling which indicated significant collateral sensitivity of the laboratory-evolved strains against VNC.

Transmission of mcr genes has been associated with an enhanced rate of COL-R emergence particularly among nosocomial isolates obtained from the ICU. [55]. However, the data on COL resistance from community isolates are still scanty. Here we examined three distinct clinical isolates of Ab for biofilm-forming and antibiotic responsiveness trade-offs. Profiling for mcr genes revealed the absence of mcr1 or mcr4 in the isolates. Quite interestingly, in one of the isolates, phenotypic attributes like cellular aggregation and biofilm formation, similar to the laboratory-evolved strains, were observed. The strains also demonstrated a similar antibiotic sensitivity profile to the evolved COL-R strains with hypersusceptibility to VNC. Additionally the study identified potentiation of CIP and fosfomycin coupled to evolution of COL-resistance.

Due to the immense genomic plasticity of Ab, even a susceptible bacterial population often manifests heteroresistance rendering the emergence of a subpopulation resistant to antibiotics [56,57]. The phenomenon is frequently reported for clinical isolates of Ab including several MDR isolates demonstrating heteroresistance against COL [58]. Identification of similar mutations in three independently evolved COL-R lines suggests the presence of a heteroresistant subpopulation within the source clonal population. Whether LPS deficiency in Ab emerges as one of the phenotypically varied subpopulations as observed in *A. baylyi* ADP1 awaits further investigation [59]. Loss of LPS due to the process of COL-resistant evolution is considered a trade-off between resistance and virulence [60]. However, the genetic regulation and systemic impact of resistance-associated genomic variations is complex and also dependent on the genomic background and selection pressure implemented. Hence, an intricate and comprehensive approach linked to multi-omics [61] might offer a more clarified view on resistance emergence-associated trade-offs and phenotypic variations.

## Supporting information

Supplemental File for Identification of Evolutionary Trade-Offs Associated with High-Level Colistin Resistance in Acinetobacter baumannii

## Acknowledgement

The authors acknowledge all the open-source software and server providers. The authors acknowledge Dr. Priyanka Bhowmik, Adamas University, and Dr. Sulagna Basu, ICMR-NIBRI for providing us with mcrt-1 and mcr-2 DNA samples and contributions to our project work. The authors are obliged to Subhash Mukhopadhyay Centre for Stem Cell Biology Regenerative Medicine, Dept. of Chemistry, and Central Instrumentation facility, Adamas University for support with instrument facilities. The authors also acknowledge Mr. Saikat Samanta, Adamas University for his constant support in laboratory activities. AB is funded by Adhoc Grant-Project ID: AMR/Adhoc/284/2022-ECD-II (ICMR, Govt. of India) and SEED grant, Adamas University.

## Availability of data and materials

The data set supporting the conclusions of this article is available in the Sequencing Read Archive (https://www.ncbi.nlm.nih.gov/sra) repository under BioProject accession no. PRJNA1188730 with sample accession numbers SRX26787849, SRX26878385, SRX26878386, and SRX26878387.

## Author contribution

Kusumita Acharya, Upasana Bhattacharya, and Shatarupa Biswas performed the experiments.

Mallika Ghosh isolated, identified, and provided the resistant clinical isolates.

Arijit Bhattacharya conceptualized the work and prepared the final manuscript.

All approved the final draft.

## Ethical approval

The study was performed with due ethical approvals from Institutional Ethical Committee, Adamas University and Institutional Biosafety Committee, Adamas University. No personally identifiable patient/ human subject information was disclosed to the researchers.

## Consent to participate

NA

## Consent for publication

NA

