## Supplemental File for Identification of Evolutionary Trade-Offs Associated with High-Level Colistin Resistance in Acinetobacter baumannii for "Identification of Evolutionary Trade-Offs Associated with High-Level Colistin Resistance in *Acinetobacter baumannii*"

Fig. S1. Schematic representation of experimental evolution of resistant mutants.

Fig. S2. Evolution of CIP-resistant *A. baumannii* mutants.

Fig. S3. Biofilm development by ciprofloxacin.

Fig. S4. Collateral sensitivity of colistin (COL)-resistant mutants against vancomycin (VNC).

Fig. S5. Fosfomycin sensitivity of colistin resistant mutants.

Table-S1. List of primers used in the study.

Table-S2. Predicted MIC vales for clinical isolates against antibiotics.

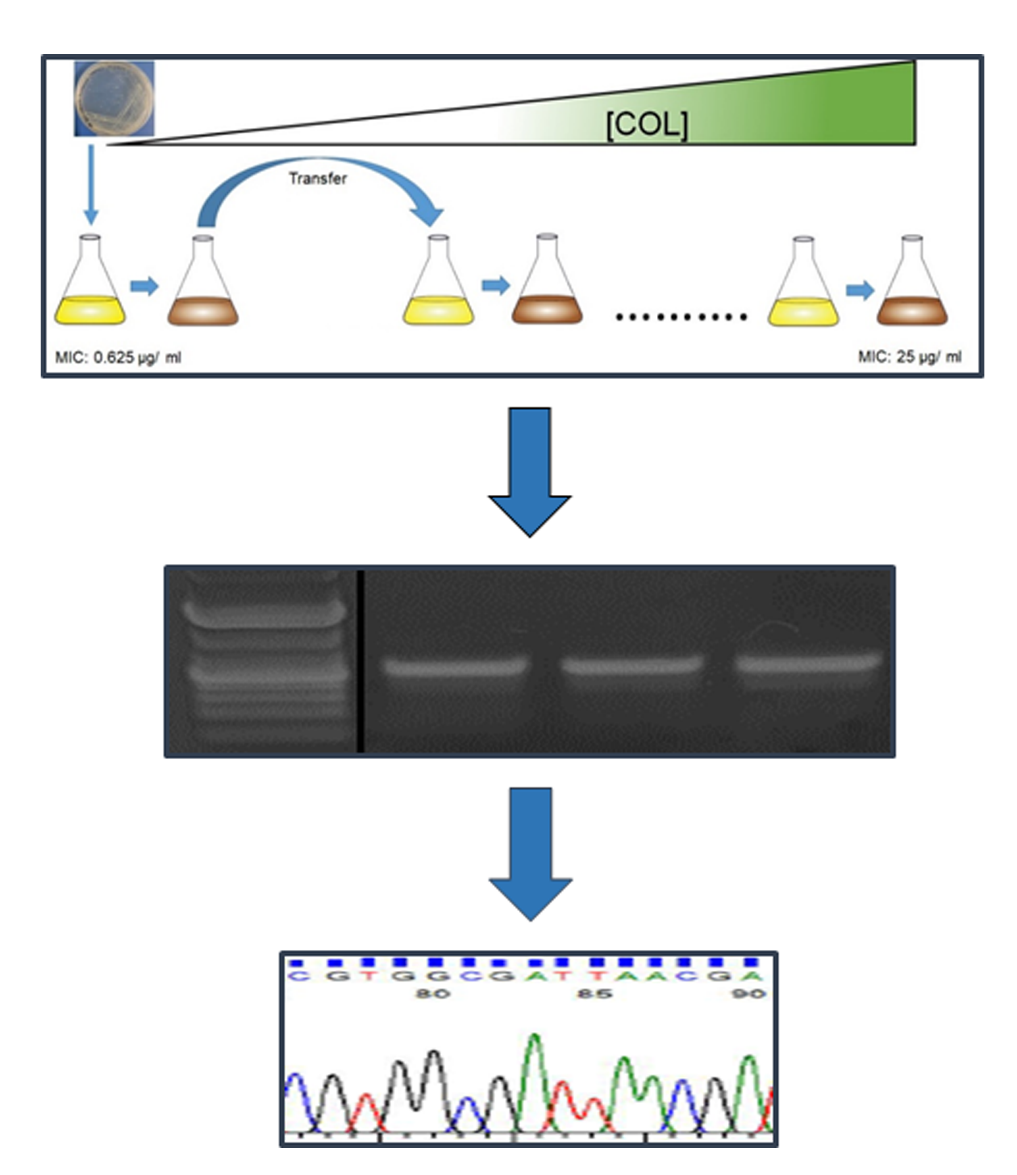

Fig. S1. Schematic representation of experimental evolution of resistant mutants. Step-wise selection for COL-resistance was performed by gradually increasing antibiotic concentration from 0.25XMIC to 20XMIC. After each round of selection the purity of the line will be confirmed by PCR with *A. baumannii* *gapdh* gene specific primer. Clones derived from the final selection was further confirmed by Sanger sequencing of the *gapdh* PCR amplicon. Similar approach was adopted for developing CIP-R mutants.

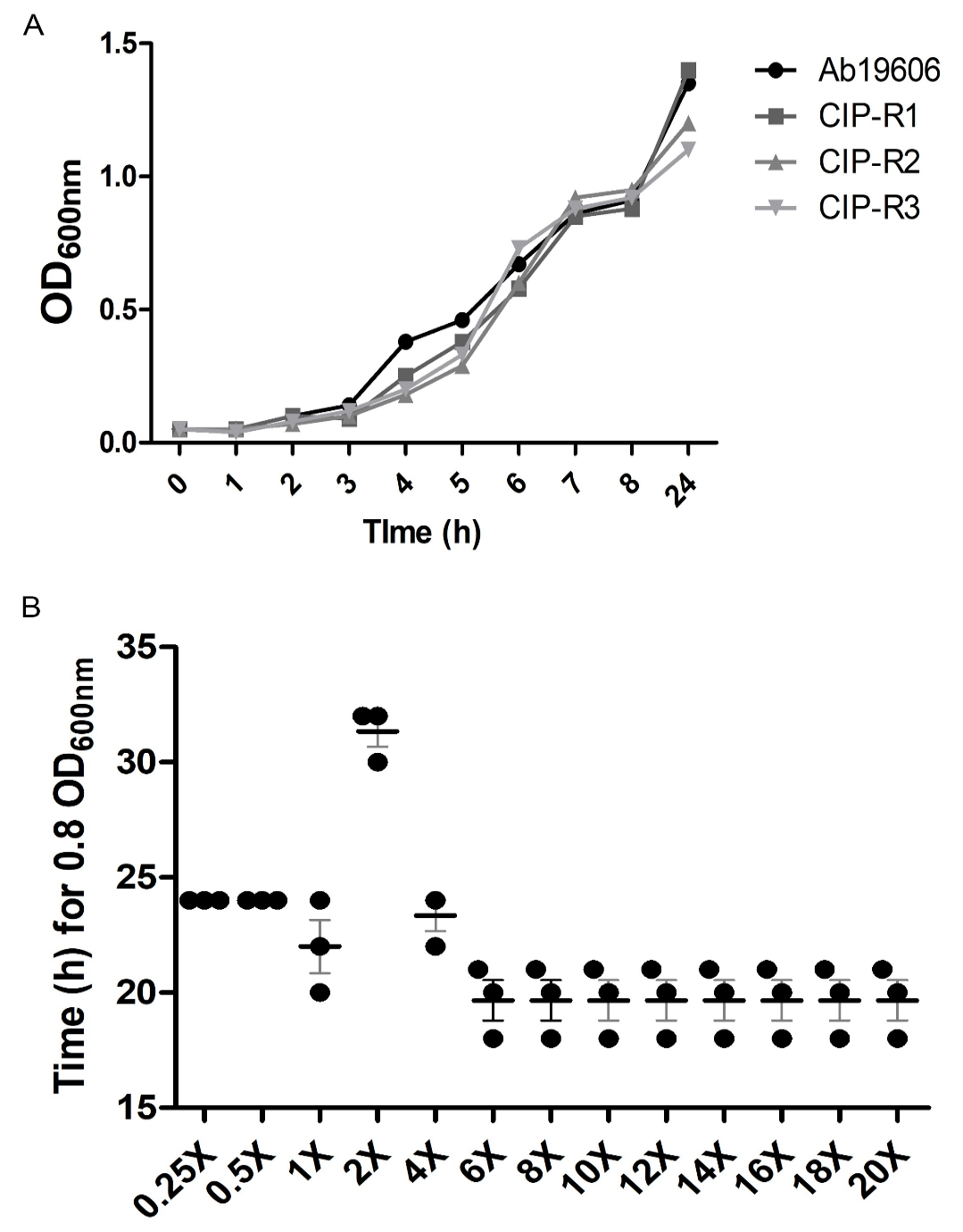

Fig. S2. Evolution of ciprofloxacin (CIP)-resistant *A. baumannii* mutants. CIP resistant lines were evolved by step-wise selection of a source clonal population against gradually increasing dose of antibiotic starting from 0.25XMIC (0.125 µg/ ml) for COL to a maximal selection at 20X MIC (10 µg/ ml). Selection dynamics (trail) at each stage of variation is projected in time needed to reach late-log phase (OD_600nm_ = 0.8) at each step of selection. Data represents mean ±SEM for three independent lines (n=3 for each) (A). (B) Growth kinetics for the Ab source, AbCOL-R1, R2, and R3 clonal populations selected against COL were performed monitoring OD_600 nm_ for 24 h while growing in absence of the antibiotic. Data represents mean ±SEM for three independent lines (n=3 for each).

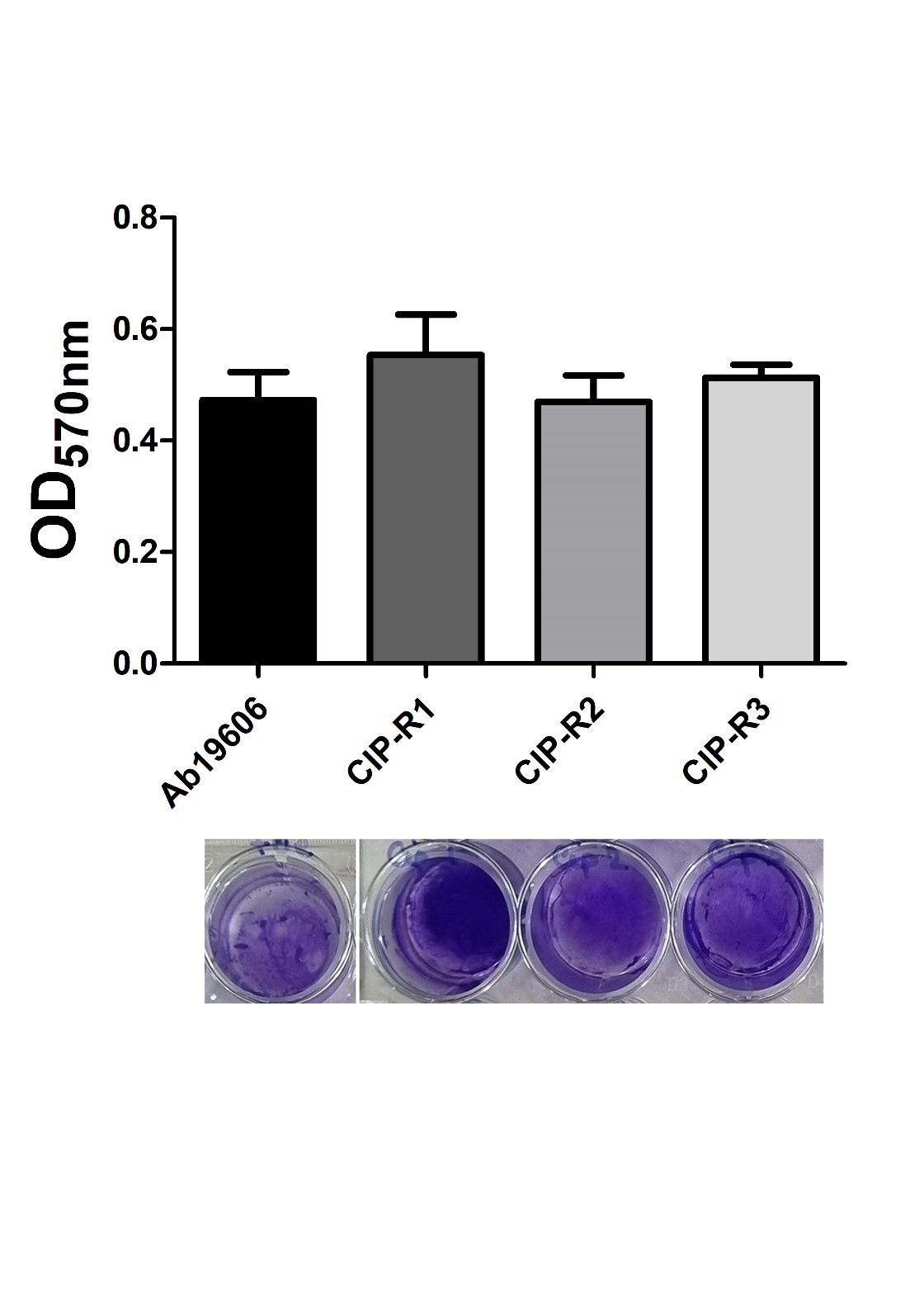

Fig. S3. Biofilm development by ciprofloxacin (CIP) resistant (CIP-R) mutants. (A) Static biofilm formation on polystyrene substratum were envisioned by crystal violate retention for each of the Ab source population (Ab19606), AbCIP-R1, R2, and R3. Results represent mean±SEM for three independent experiments. ***p<0.001, two tailed paired Student t-test. In set: representative data from one replicate of the experiment demonstrating biofilm development.

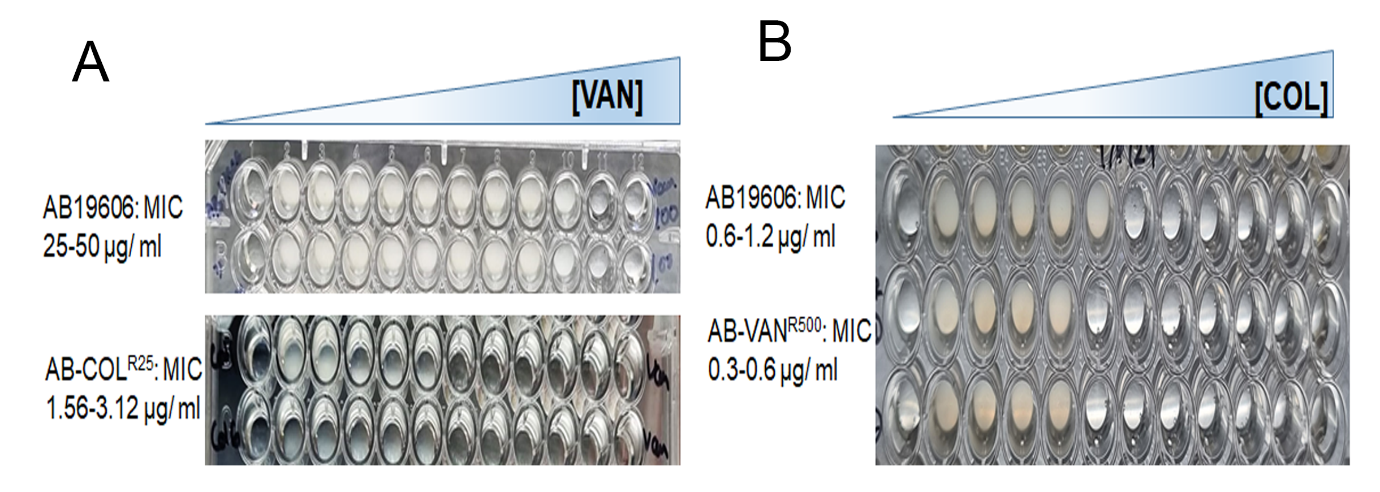

Fig. S4. Collateral sensitivity of colistin (COL)-resistant mutants against vancomycin (VNC). MIC were determined for COL-resistant lines (AbCOL-R1, R2, and R3) and the source clonal population of Ab19606 cultured in parallel while generating resistant lines against VNC. Data from a representative replicate is shown and the mean value for the lined are mentioned separately.

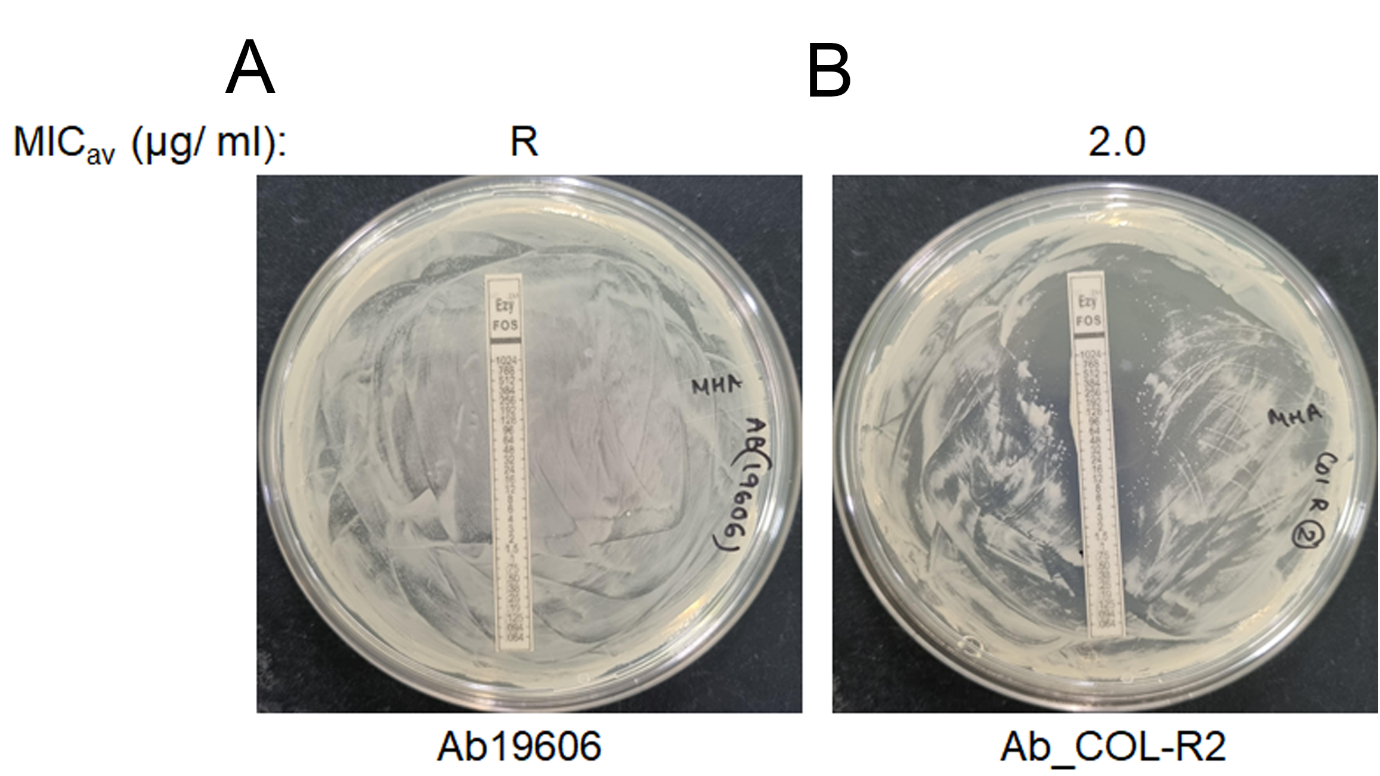

Fig. S5. Fosfomycin sensitivity of colistin resistant mutants. Collateral sensitivity against fosfomycin was assessed using E-strip MIC test against Ab19606 source clonal population (A) and AbCOL-R clonal populations. A representative plate for AbCOL-R2 is shown (B).

Table-S1. List of primers used in the study. The primers used in the study are enlisted. Sequences are depicted in 5’ to 3’ direction.

| Primer_ID | Sequence (5'-3') |
| --- | --- |
| 16s_8F | agagtttgatcctggctcag |
| 16s_1492R | ggttaccttgttacgactt |
| Abgapdh_F | atgcaacgtatcgccatt |
| Abgapdh_R | tcgtacatgacacactcgat |
| mcr1_sbmx_320_F | agtccgtttgttcttgtggc |
| mcr1_sbmx_320_R | agatccttggtctcggcttg |
| mcr2_sbmx_715_F | caagtgtgttggtcgcagtt |
| mcr2_sbmx_715_R | tctagcccgacaagcatacc |
| mcr3_sbmx_929_F | aaataaaaattgttccgcttatg |
| mcr3_sbmx_929_R | aatggagatccccgttttt |
| mcr4_sbmx_1116_F | tcactttcatcactgcgttg |
| mcr4_sbmx_1116_R | ttggtccatgactaccaatg |
| mcr5_sbmx_1644_F | atgcggttgtctgcatttatc |
| mcr5_sbmx_1644_R | tcattgtggttgtccttttctg |
| mcr6_sbmx_260_F | atttagtagtcactacgccagtttc |
| mcr6_sbmx_260_R | cacgactatagccattaaactgcac |
| mcr7_sbmx_390_F | tcgttatctcggttcctctggt |
| mcr7_sbmx_390_R | cgtaatcctgatagtaaagtgcgg |
| mcr8_sbmx_582_F | ttattccatatttatgcaggcctc |
| mcr8_sbmx_582_R | aaagataggggttggttacccg |
| mcr9_sbmx_852_F | ggccgcaataactcgacatt |
| mcr9_sbmx_852_R | cagatatagcccgctttcgc |
| mcr10_sb_1387_F | tgcagcgcttaactttgtgt |
| mcr10_sb_1387_R | gcagatacagtccgctctct |
| Ab_PmrB1_F | ggcaatcataatttaaaatttcggg |
| Ab_PmrB1_R | cacgctcttgtttcatttaaatgt |
| Ab_PmrB2_F | ggcaatcataatttaaaatttcggg |
| Ab_PmrB2_R | gcatcggcaataaattgcttc |
| Ab_PmrB3_F | gaagcaatttattgccgatgc |
| Ab_PmrB3_R | cacgctcttgtttcatttaaatgt |
| Ab_PmrB1_qRT_F | gaagtacatgattatcctcaagagc |
| Ab_PmrB1_qRT_R | tacgtgccaaacctttgctt |
| pmrA_seq_F | atgacaaaaatcttgatgattgaag |
| pmrA_seq_R | ttatgattgccccaaacggt |
| pmrA_qrt_F | tctgctcgagatcaattacaaaacc |
| pmrA_qrt_R | cgtcgcaatatgctgttcaa |
| lpxA_seq_F | atgagcaatcacgatttaatcca |
| lpxA_seq_R | ttagcgcacaattccacg |
| lpxA_qrt_F | attcgcgaacattgcagctt |
| lpxA_qrt_R | accacctacaataacgtgatcacc |
| lpxC_seq_F | atggtgaaacagcgtactctcaa |
| lpxC_seq_R | ttatgtcacactcacgtatggaa |
| lpxC_qrt_F | tggcctgcgtgaacaagat |
| lpxC_qrt_R | gttgcagactgatattctttggc |

Table-S2. Predicted MIC vales for clinical isolates against antibiotics. MIC value as predicted by automated AST assay with VITEK 2 system for *A. baumannii* DLPL20, DLPL37, and DLPL47 are mentioned.

| **Antibiotics** | **MIC (µg/ ml)** | | |
| --- | --- | --- | --- |
|  | **DLPL20** | **DLPL37** | **DLPL47** |
| Piperacillin/Tazobactum | >=128 | >=129 | <=4 |
| Ceftazidime | 32 | 2 | <=0.12 |
| Cefoperazone/Sulbactam | 32 | <=8 | <=8 |
| Cefepime | 16 | 0.5 | <=0.12 |
| Imipenem | 2 | <=0.5 | <=0.5 |
| Meropenem | 8 | <=0.25 | <=0.25 |
| Amikacin | >=64 | 4 | 16 |
| Gentamicin | >=16 | <=1 | <=1 |
| Ciprofloxacin | >=4 | 0.5 | <=0.06 |
| Levofloxacin | >=8 | <=0.12 | <=0.12 |
| Minocyclin | 4 | <=0.5 | 4 |
| Colistin | >=16 | 8 | >=16 |
| Trimethoprim/Sulfamethoxazole | <=20 | <=20 | <=20 |
